# Individual trajectories for recovery of neocortical activity in disorders of consciousness

**DOI:** 10.1101/2024.03.11.584433

**Authors:** Prejaas K.B. Tewarie, Romesh Abeysuriya, Rajanikant Panda, Pablo Nùñez, Marie M. Vitello, Glenn van der Lande, Olivia Gosseries, Aurore Thibaut, Steven Laureys, Gustavo Deco, Jitka Annen

**Author notes:** corresponding author: Prejaas K.B. Tewarie. These authors contributed equally to this work (Gustavo Deco, Jitka Annen).

## Abstract

The evolution from disturbed brain activity to physiological brain rhythms can precede recovery in patients with disorders of consciousness (DoC). Accordingly, intriguing questions arise: What are the pathophysiological factors responsible for disrupted brain rhythms in patients with DoC, and are there potential pathways for individual patients with DoC to return to normal brain rhythms? We addressed these questions at the individual subject level using biophysical simulations based on electroencephalography (EEG). The main findings are that unconscious patients exhibit a loss of excitatory corticothalamic synaptic strength. Synaptic plasticity in this excitatory corticothalamic circuitry fosters physiological brain rhythms in the selection of patients with DoC. The extent to which this occurred was correlated with cerebral glucose uptake. The current findings emphasize the importance of excitatory thalamocortical activity in reestablishing normal brain rhythms after brain injury and show that biophysical modelling of the corticothalamic circuitry could help select patients that might be potentially receptive to treatment and undergo plasticity.

## Introduction

Disorders of consciousness (DoC) occur in a small proportion of comatose survivors with severe brain injury. While the etiology varies among patients, common etiologies include severe traumatic brain injury (TBI), stroke, and cardiac arrest. Irrespective of the etiology, these patients can be grouped into the unresponsive wakefulness syndrome (UWS), characterized by the presence of eye-opening and reflexive behaviors, and the minimally conscious state (MCS), characterized by inconsistent conscious behaviors, such as command following or visual pursuit. There is widespread reduction of cerebral metabolism, commonly measured using [^18^F]Fluorodeoxyglucose Positron Emission Tomography (FDG-PET), and disrupted neocortical activity in DoC due to damaged neuronal circuitry at the cellular level J. T. Giacino, Fins, et al. 2014. Restoration of cerebral activity (e.g., measured with electroencephalography (EEG) or FDG-PET) is believed to precede clinical recovery Bareham et al. 2020; Thibaut, Panda, et al. 2021. Patients in whom preservation of cerebral activity was observed in the absence of behavior were referred to as MCS stars (MCS^*^) Gosseries, Zasler, and Laureys 2014. Hence, exploring potential routes for the restoration of cerebral activity is crucial for understanding clinical recovery in patients with DoC.

Neocortical activity is commonly measured by EEG. This activity represents the electrical signals generated by neural populations under the skull, orchestrated by corticocortical and thalamocortical connections. EEG signals in healthy subjects can usually be decomposed into an aperiodic component (1/f component) and a periodic component or spectral peaks Donoghue et al. 2020. A proposed qualitative model for the evolution of the EEG power spectrum in DoC is often referred to as the “ABCD” model, derived from the mesocircuit hypothesis Edlow, Claassen, et al. 2021. This model states that neocortical activity in severe brain damage leading to UWS is restricted to the aperiodic part, which is dominated by a high delta power (i.e., ¡ 4 Hz). Recovery of consciousness is believed to co-occur with the emergence of a spectral peak in the lower frequency range (around 7 Hz), with a shift towards higher frequencies (first around 10 Hz and potentially around 20 Hz) during further recovery Edlow, Claassen, et al. 2021. The relevance of EEG measurements in DoC is underscored by the notion that the re-emergence of physiological EEG features is associated with behavioral recovery Colombo et al. 2023; Jørgensen and Holm 1998; Maschke et al. 2023; Thibaut, Panda, et al. 2021 and glucose uptake Annen et al. 2023.

Several hypotheses may explain the recovery of the EEG power spectrum in patients with DoC. One of these hypotheses, the meso-circuit hypothesis, states that the return of spectral peaks in the EEG power spectrum is a result of the restoration of excitation in the thalamocortical circuit and, more specifically, a more widespread cortico-striato-pallido-thalamic network N. D. Schiff 2010. This hypothesis also postulates widespread deactivation of excitatory synaptic activity across the cerebral cortex, resulting in a global hyperpolarized state in DoC, which is known as ”disfacilitation” Edlow, Claassen, et al. 2021. Recovery in this situation would require boosting the excitation to induce a shift from this hyperpolarized state. Restoration of excitatory processes presumably depends on neuronal repair and synaptic plasticity Colombo et al. 2023, and both are assumed to occur in DoC to some extent Werner and Stevens 2015. The potential roles of neuroprotective and neurostimulating drugs that induce cerebral plasticity, resulting in partial recovery of consciousness, underscore this view Demertzi et al. 2011.

The mechanisms underlying the recovery of EEG power spectra in patients with DoC cannot be directly inferred from the EEG data. However, using biophysical models of macroscopic brain activity, we cannot only infer biophysical model parameters (e.g., synaptic properties not measured empirically with EEG) from EEG power spectra of individual patients but also model subject-specific synaptic plasticity R. Abeysuriya and P. Robinson 2016. Our biophysical model includes excitatory and inhibitory intracortical, intrathalamic, and corticothalamic synaptic strengths as well as synaptic time constants and axonal conduction delays. This model allowed us to test the roles of excitation and inhibition in thalamocortical synaptic connections. It allows the identification of the circuits that are most affected in patients with DoC, for example, thalamocortical or intracortical circuits. This allowed us to model the subject-specific plasticity of synapses and their effects on brain rhythms. Well-known types of plasticity commonly used in corticothalamic biophysical models are Hebbian and homeostatic plasticities Romesh G Abeysuriya et al. 2018; Fung, Haber, and P. Robinson 2013; Magee and Grienberger 2020. The former is a positive feedback-mediated form of plasticity in which synapses between presynaptic and postsynaptic neurons that are coincidently active are strengthened. The latter is a negative feedback-mediated form of plasticity, also known as synaptic scaling, which maintains network activity at the initial level.

In the current work, we will address three questions: 1) Which synaptic parameter determines disrupted EEG in patients with DoC the most? 2) What are the potential cortical or corticothalamic routes of plasticity that lead to neurophysiological recovery in individual patients with DoC? 3) To what extent do the routes for neurophysiological recovery relate to metabolic preservation as potential biomarkers of plasticity?

## Results

### Estimating corticothalamic model parameters for disturbed EEG patterns in patients with DoC

The power spectra for each subject were estimated and averaged across (occipital, temporal and parietal) electrodes, resulting in a single power spectrum per subject. We used a Markov Chain random walk to estimate the parameters of the biophysical corticothalamic model R. Abeysuriya and P. Robinson 2016. This corticothalamic model includes a cortical excitatory and cortical inhibitory population, a thalamic relay, and a reticular population. The postsynaptic membrane potential of a population is modulated by the firing activity of the presynaptic populations. The impact of presynaptic input on the postsynaptic membrane potential also depends on the mean number of synapses between the presynaptic b and postsynaptic population a (modelled as synaptic strength *ν*_*ab*_) and on the closing and opening rates of synaptic channels characterized by the synaptic decay and rise constants (*α* and *β*). The average postsynaptic membrane potential is transformed into firing activity in the cell bodies of a population. This process results in activity propagation in a closed loop between the thalamic and cortical populations, where the firing rate propagated between the thalamus and cortex is delayed by *t*_0_. The synaptic parameters *ν*_*ab*_ are transformed in a linear version of the model to the gain parameters *G*_*ab*_ and closed loop parameters *G*_*aba*_ = *G*_*ab*_*G*_*ba*_. Given this transformation, the gain parameters did not strictly indicate the equivalence of synaptic strengths.

Figure 2A shows the empirical power spectra (red) and model-estimated power spectra (blue) averaged across the subjects within each group. Power spectra from patients with UWS are mostly characterized by an aperiodic component without a spectral peak, whereas power spectra from patients with MCS are predominantly characterized by an aperiodic component with a subtle peak in the theta-band (4-8 Hz). The fraction of patients with a spectral peak in the (lower) frequency bands was 0.21 for UWS and 0.44 for MCS (p = 0.026, *χ* = 4.97). The power spectrum of healthy control subjects was characterized by an aperiodic component (1/f part) and a spectral peak in the alpha band at approximately 10 Hz. For all groups, the model-estimated power spectra (blue curves) provide a close approximation to the empirical power spectra, as evident from Fig. 2A, both visually and in terms of the goodness-of-fit (GOF) in the rightmost panel. There was no significant difference in GOF between the groups (p *>* 0.05). Figures S2–S4 show the estimated and empirical power spectra for the individual patients. This indicates that the individual fits only captured the global shape of the power spectra, including the most prominent peaks. This may lead to the omission of details in the estimation spectra but avoids overfitting.

**Figure 1:**
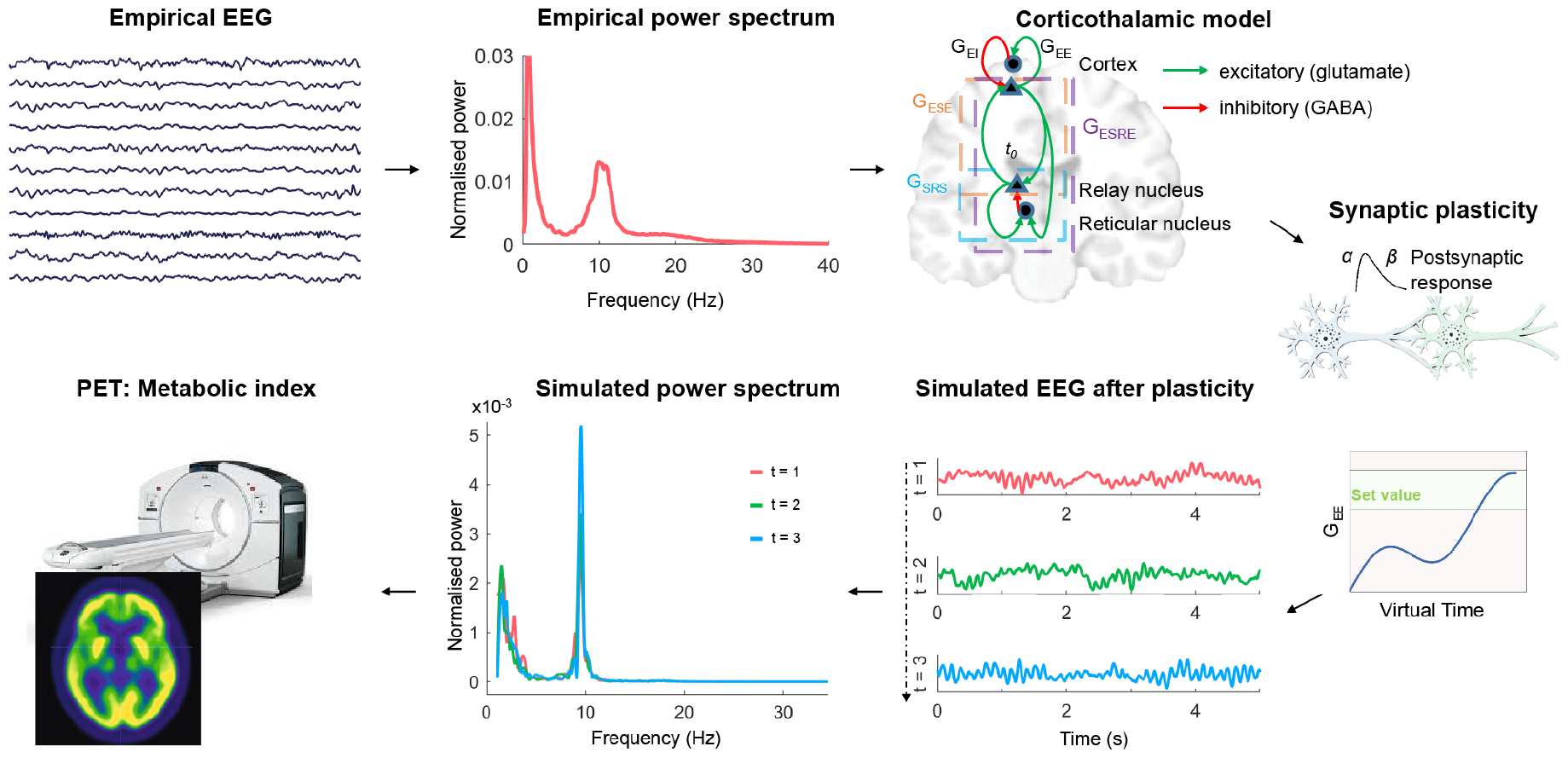
Overview of the pipeline. The first part of the analysis entails the estimation of biophysical model parameters from individual EEG data. We use a Markov Chain random walk to estimate biophysical parameters from subject-specific whole-brain power spectra. This is followed by subject-specific simulations using either corticothalamic or cortical plasticity. Potential links with available FDG-PET data are further analysed. Abbreviations: excitatory corticothalamic synaptic strengths (*G*_*ESE*_), inhibitory corticothalamic synaptic strengths (*G*_*ESRE*_), Excitatory cortical synaptic 158 strengths (*G*_*EE*_), inhibitory cortical synaptic strengths (*G*_*EI*_), intrathalamic synaptic strengths (*G*_*SRS*_), synaptic decay and rise constants (*α* and *β*), corticothalamic time delay (*t*_0_), electroencephalography (EEG).

**Figure 2:**
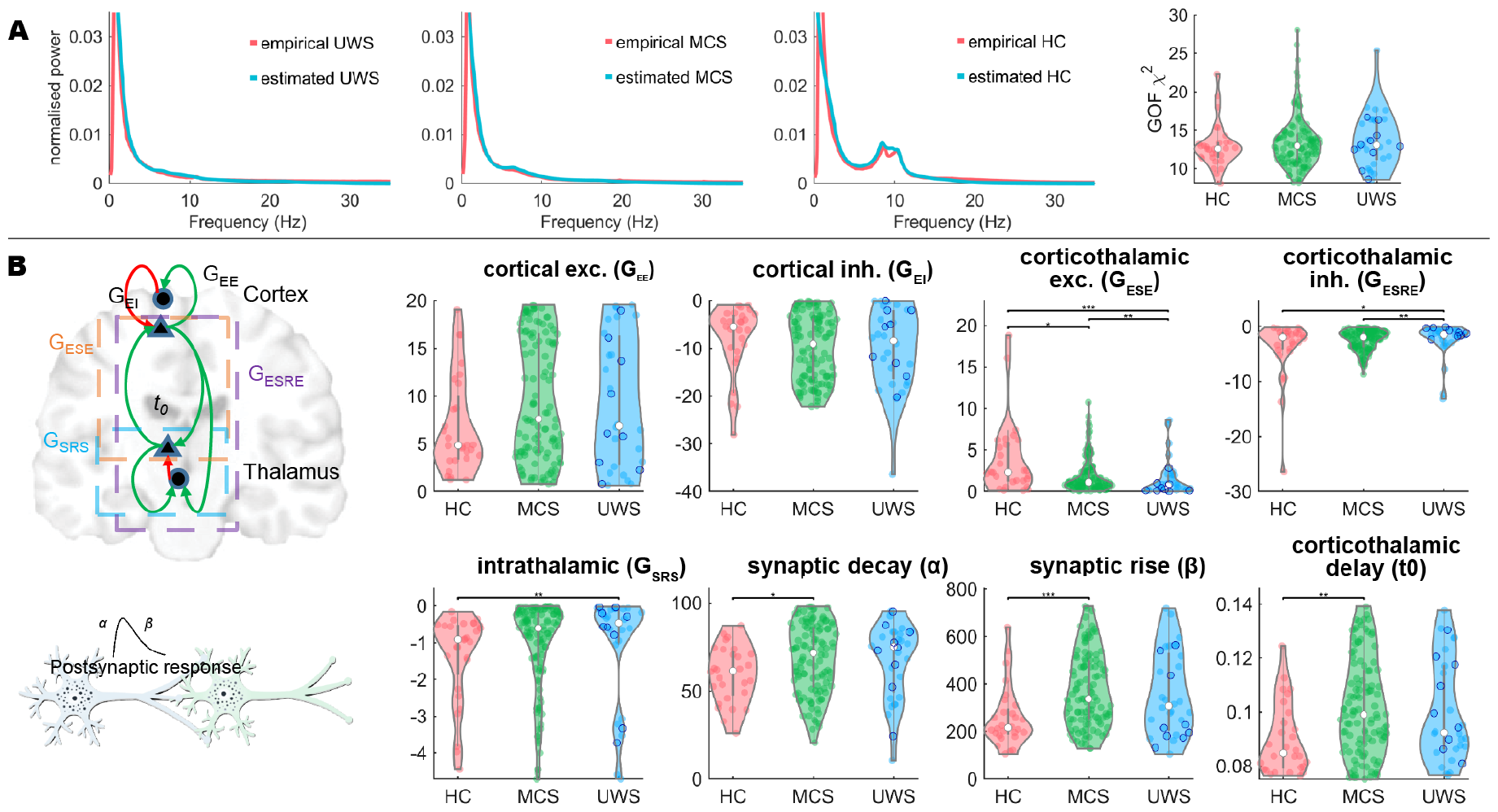
Estimating corticothalamic model parameters in DoC. Panel A shows the group-averaged empirical power spectra and model-estimated power spectra for the different groups alongside the goodness of fit of the estimation for the different groups. Panel B shows a graphical description of the model parameters and their estimates in the different groups. Blue circles in the UWS group denote MCS^*^ patients. Both excitatory and inhibitory corticothalamic synaptic strengths differ between MCS and UWS. Abbreviations: healthy controls (HC), minimal conscious state (MCS), unresponsive wakefulness syndrome (UWS), excitatory corticothalamic synaptic strengths (*G*_*ESE*_), inhibitory corticothalamic synaptic strengths (*G*_*ESRE*_), Excitatory cortical synaptic strengths (*G*_*EE*_), inhibitory cortical synaptic strengths (*G*_*EI*_), intrathalamic synaptic strengths (*G*_*SRS*_), synaptic decay and rise constants (*α* and *β*), corticothalamic time delay (*t*_0_). *\** = *p <* 0.05, *\*\** = *p <* 0.01, *\* * \** = *p <* 0.001. (FDR corrected)

The groups did not differ in cortical excitatory (*G*_*ee*_) or cortical inhibitory (*G*_*ei*_) synaptic gains (Fig. 2B). However, all groups differed in terms of excitatory corticothalamic synaptic gains (*G*_*ese*_) and inhibitory corticothalamic synaptic gains (*G*_*esre*_). In particular, the excitatory corticothalamic synaptic gain (*G*_*ese*_) could differentiate between the MCS and UWS groups. Patients in the UWS group were characterized by a smaller excitation and larger inhibition of the thalamocortical system. There was a loss of inhibition in the intrathalamic loop in DoC (*G*_*srs*_); however, this could not differentiate between UWS and MCS. Furthermore, there were considerably longer synaptic decay and rise constants (*α* and *β*) for the MCS group compared to the healthy control (HC) group, although the UWS group did not differ from either the HC or MCS group. The same holds true for the corticothalamic time delay *t*_0_ with longer time delays in the MCS group than in the healthy control group, and no differences between the MCS and UWS groups were observed. Of note, the aforementioned significant differences between the groups could not be explained merely by etiology (see Fig. S4).

### Potential routes for recovery of EEG activity in individual patients with DoC

We studied two potential mechanisms for the recovery of EEG activity in patients with DoC: 1) the role of excitatory synaptic plasticity in the cortex and 2) the role of excitatory synaptic plasticity in the corticothalamic loop. We employed the full nonlinear model in this context and simulated model Equations 1–3 (see Materials and Methods) with subject-specific model parameters obtained from the previous analysis (Fig. 2B). The synaptic parameters *G*_*ab*_ translate to *ν*_*ab*_ in the fully nonlinear model regime, and we applied synaptic plasticity (Equation 14) to the excitatory corticothalamic loop *ν*_*es*_ and the excitatory intracortical loop *ν*_*ee*_.

Figure 3A shows examples of the subject-specific simulations of corticothalamic plasticity. The red line shows the empirical power spectrum superimposed on several power spectra during the evolution of synaptic plasticity (transformation from red to green). The effects of corticothalamic synaptic plasticity differed between the subjects. While some subjects developed a spectral peak in the theta band or even the alpha band, other subjects did not develop a clear spectral peak despite the induction of corticothalamic synaptic plasticity. For example, a patient with UWS due to anoxia did not show an apparent spectral peak after corticothalamic plasticity, whereas a patient with MCS due to TBI developed an evident alpha peak only after the corticothalamic plasticity was set (Fig. 3A). The implementation of cortical plasticity alone produced a different picture (Fig. 3B). Although cortical synaptic plasticity led to a different slope of the power spectrum, no spectral peak was induced by cortical plasticity alone in any of the patients. Figure 3B shows the power spectra for the same groups demonstrated in Fig. 3A.

**Figure 3:**
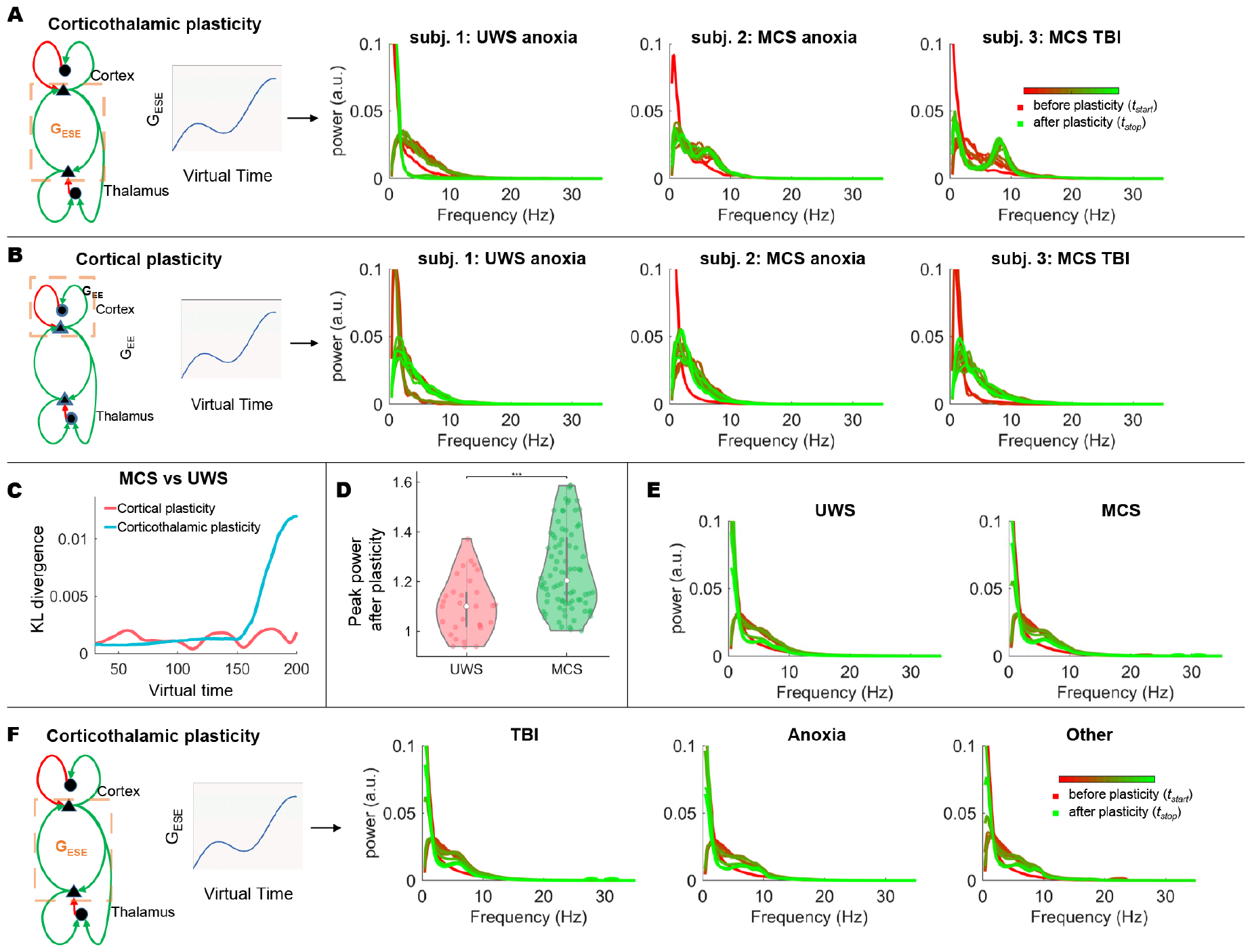
Modelling synaptic plasticity in DoC. Panels A and B show simulated power spectra for individual subjects during excitatory corticothalamic (A) and excitatory cortical synaptic plasticity (B). Panels D and E show the Kullback-Leibler (abbreviated to KL) divergence during either cortical plasticity (D) or corticothalamic plasticity (E). The power corresponding to the peak frequency after completion of corticothalamic plasticity is illustrated for the three different groups (UWS, MCS, and EMCS), with a significant difference (*p <* 0.001) denoted by ^***^. Panel G shows the group-averaged power spectra during the course of corticothalamic plasticity for the UWS, MCS, and EMCS groups. The same group-averaged power spectra are illustrated for different etiologies (H). Abbreviations: healthy controls (HC), minimal conscious state (MCS), unresponsive wakefulness syndrome (UWS), excitatory corticothalamic synaptic strengths (*G*_*ESE*_), inhibitory corticothalamic synaptic strengths (*G*_*ESRE*_), Excitatory cortical synaptic strengths (*G*_*EE*_), inhibitory cortical synaptic strengths (*G*_*EI*_), intrathalamic synaptic strengths (*G*_*SRS*_), synaptic decay and rise constants (*α* and *β*), corticothalamic time delay (*t*_0_).

The results of the individual spectra suggest that spectral peaks are more likely to occur during the progression of synaptic corticothalamic plasticity in patients with MCS than in those with UWS. The fraction of patients that showed a clear peak after corticothalamic plasticity that was not present in the initial and original empirical data was 0.2 for UWS and 0.45 for MCS (p = 0.019, *χ* = 5.43). This was also assessed using the Kullback-Leibler divergence, which was used to quantify the difference in the shape of the power spectrum between the UWS and MCS groups during plasticity. First, regarding cortical plasticity, we observed that the Kullback-Leibler divergence remained approximately constant during the effectuation of synaptic plasticity (Fig. 3C). Second, for corticothalamic plasticity, we observed that the power spectra between the UWS and MCS groups diverged as plasticity advanced (Fig. 3C). This effect was probably driven by a significantly stronger occurrence of a spectral peak in the higher levels of consciousness than in the lower levels of consciousness (Fig. 3D), which could also be observed from the group-averaged power spectra in the respective groups after corticothalamic plasticity (Fig. 3E). Finally, we analyzed the effect of corticothalamic plasticity in patients with different etiologies, as the contributions of certain etiologies may differ between the MCS and UWS groups (Fig. 3F). The group-averaged power spectra showed that the reoccurrence of a spectral peak might be more prominent in the TBI subgroup than in the anoxic subgroup (Fig. 3F).

### Neurophysiological correlates of FDG-PET findings in DoC

To better characterize the relationship between cerebral integrity and (the ability to recover a more normal) peak frequency of power spectral density, we associated the peak frequency with metabolic activity. We extracted the peak frequency of the empirical power spectrum for all the participants for whom a peak was present before the application of plasticity (UWS, n = 7 of 36; MCS, n = 42 of 95). This was also performed for the power spectra extracted from the simulated EEG data after corticothalamic plasticity (UWS, n = 13 of 36; MCS, n = 80 of 95). The scatter plots in Fig. 4A show the relationship (or lack thereof) between the metabolic index of the best-preserved hemisphere and peak frequency from the empirical power spectra. A strong positive correlation between these two entities was observed in the UWS group, whereas no significant correlation was observed in the MCS group. The correlation in the UWS group was mainly driven by the presence of patients with MCS^*^ (green in Fig. 4). However, for the modelled power spectra obtained after corticothalamic plasticity, we observed strong to moderate correlations with the metabolic index in both groups (UWS and MCS).

**Figure 4:**
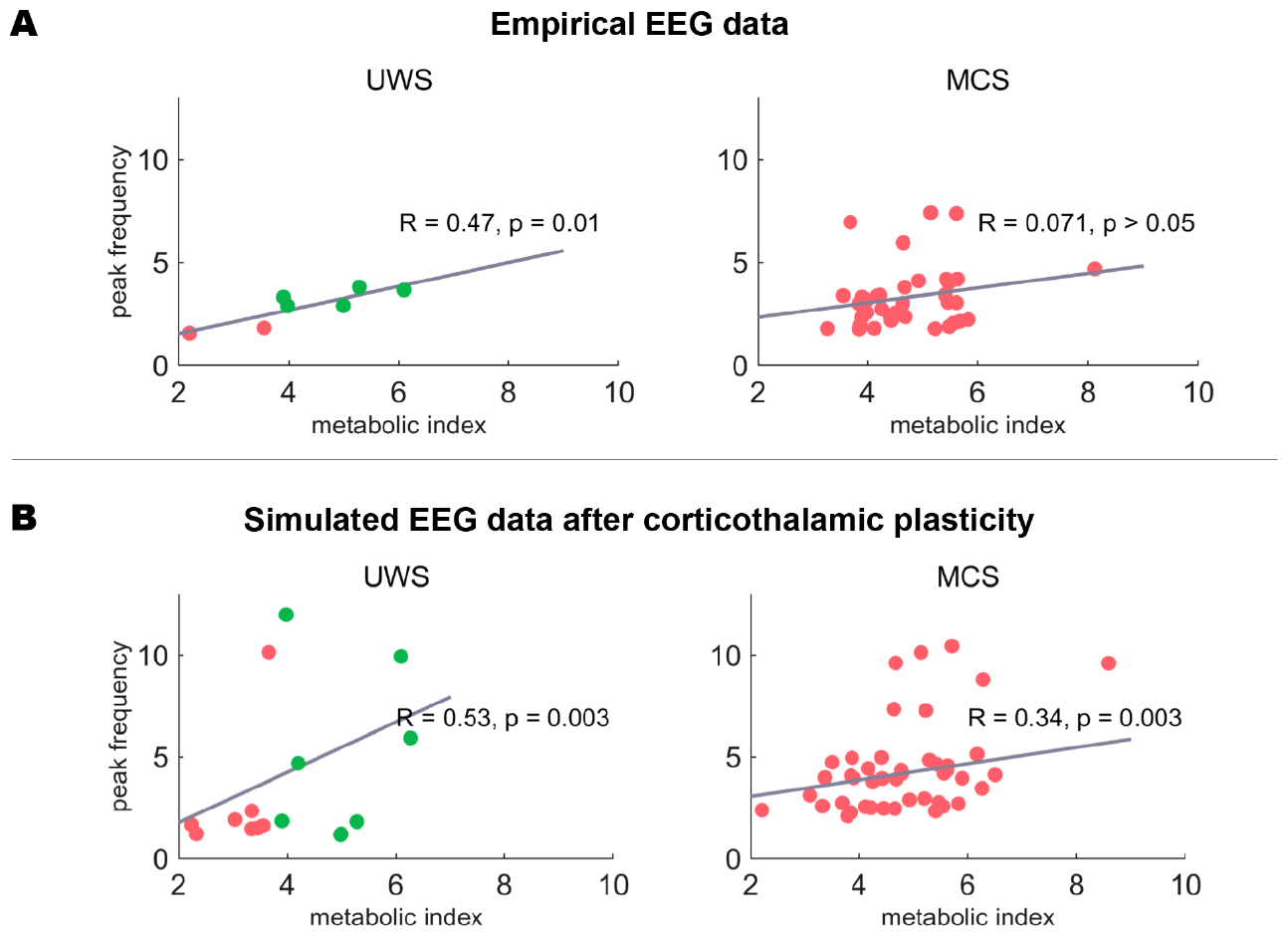
The relationship between modelled synaptic plasticity and FDG-PET findings in DoC. Panel A shows scatter plots and correlations between the peak frequencies of empirical EEG data with the metabolic index from FDG-PET. Panel B shows scatter plots and correlations between the peak frequency of simulated EEG signals after corticothalamic plasticity and the metabolic index from FDG-PET. For the modelled peak frequency, significant correlations in all groups were observed, whereas for the empirical peak frequency, a significant correlation for only the UWS group was observed. Green circles in the UWS group denote MCS^*^ patients.

## Discussion

Clinical recovery in patients with DoC may be dependent on sufficient cerebral metabolism and the recovery of physiological brain rhythms, as measured by EEG (6). Insight into the evolution of physiological EEG rhythms in DoC requires knowledge of the most important pathophysiological determinants of disturbed EEG activity. Using parameter estimation, we demonstrated that disturbed brain rhythms in DoC were likely the result of delayed propagation of neuronal activity between the thalamus and cortex, delayed synaptic responses, and loss of synaptic strength within the thalamus, especially loss of synaptic strength between the thalamus and cortex. Corticothalamic synaptic strength was not only lower in patients with DoC but also sensitive in differentiating between patients with MCS and UWS. The important role of corticothalamic synapses was strengthened by the observation that plasticity in these synapses specifically resulted in the recovery of physiological brain rhythms (e.g., theta and alpha rhythms) indicative of thalamocortical modulation in some patients. The extent to which this occurred correlated with whole-brain metabolic activity as measured by FDG-PET. Recovery of theta and alpha brain rhythms cannot be achieved merely by cortical synaptic plasticity.

The current findings are in agreement with the mesocircuit hypothesis, which states that the slow EEG activity observed in patients with DoC results from the quiescent activity of the central thalamus and complete deafferentiation of excitatory corticothalamic synapses (6). The mesocircuit model predicts that the power spectrum in DoC recovers its shape from a 1/f spectrum to a 1/f spectrum superimposed by peaks in the theta and alpha bands. While theta activity may emerge only from cortical neurons, alpha activity may emerge from increased activity in the central thalamus and the restoration of corticothalamic synaptic connectivity Forgacs et al. 2017. However, to date, this prediction has mostly been verified indirectly using structural MRI Lutkenhoff et al. 2015, fMRI Coulborn et al. 2021; Panda, Thibaut, et al. 2022, FDG-PET Fridman et al. 2014, or in vivo animal studies Timofeev et al. 2000. The current findings reveal a more direct relationship between qualitative model predictions of the mesocircuit hypothesis and EEG findings in DoC. We showed that disturbed EEG in patients with DoC may be the result of both axonal and synaptic damage, especially damage to the excitatory corticothalamic synapses. The latter was also sensitive in distinguishing between MCS and UWS. Another observation in agreement with the mesocircuit hypothesis was that intrathalamic synaptic strength was weaker in patients with DoC than in healthy controls, which could lead to reduced excitatory output towards the cortex.

The presumed role of excitatory corticothalamic synapses in the recovery of EEG rhythms in DoC was further strengthened by the finding that the plasticity of corticothalamic synapses, rather than merely cortical synapses, could result in the re-emergence of the 1/f spectrum superimposed by peaks in some patients. Interestingly, the extent to which the spectral peaks re-emerged was related to the metabolic index extracted from FDG-PET. The relationship between the modelled spectral peak and the metabolic index was stronger than that between the patients’ empirical spectral peak and their metabolic index. The relationship between the occurrence of spectral peaks on EEG after corticothalamic synaptic plasticity and the metabolic index suggests that the metabolic index from FDG-PET can encompass information about the potential to recover and decode the residual capacity of the thalamocortical system in the brain Thibaut, Panda, et al. 2021. This suggests that the functional integrity of neurons, as measured using glucose, is a good index of potential local and global network interactions that give rise to higher peak frequencies. This is in agreement with a previous functional MRI study suggesting that stronger connectivity (albeit purely cortical networks) is related to a higher metabolic index in patients with DoC Di Perri et al. 2016; Panda, López-González, et al. 2023. Furthermore, we observed that plasticity induced a spectral peak in a subset of patients. High response rates are biologically unrealistic in patients with severe brain damage, some of whom may never recover. Indeed, the number of patients with anoxic brain injury leading to global atrophy and recovery is lower than the number of patients with local traumatic brain injury Whyte et al. 2009. In line with these observations regarding spontaneous recovery, we found that patients with a traumatic brain injury had a higher probability of recovering a spectral peak than patients with anoxic brain injury. Finally, we observed that the proportion of patients in whom plasticity altered the power spectra aligns with treatment response rates reported in empirical studies reviewed by Thibaut et al. Thibaut, N. Schiff, et al. 2019. These important insights increase the biological plausibility and clinical relevance of this model.

This study has several potential clinical implications. As corticothalamic plasticity induces faster alpha oscillations in some patients, while failing to induce alpha oscillations in others, it could identify which subjects could be sensitive to (noninvasive) interventions in DoC Edlow, Sanz, et al. 2021, such as amantadine or transcranial direct current stimulation Martens et al. 2020; Thibaut, N. Schiff, et al. 2019; Thibaut, Wannez, et al. 2017. The results suggest that in some patients, overall synaptic and axonal damage is too severe, and promoting corticothalamic plasticity in these patients is insufficient to return to physiological brain rhythms. In some patients, there may be a residual capacity of the brain that can be assessed by promoting corticothalamic plasticity. For example, the results of deep brain stimulation of the central thalamus in patients with DoC have been mixed Vanhoecke and Hariz 2017, potentially because of the suboptimal selection of patients with sufficient residual capacity, despite the observation that plasticity may be promoted by deep brain stimulation in other clinical populations Van Hartevelt et al. 2014. Recent work has shown that low-frequency stimulation of the centromedian-parafascicular complex (CM-Pf) in the thalamus in DoC could promote the regeneration of whole-brain communication in the alpha band Arnts et al. 2022. Hence, there may be a role for subject-specific corticothalamic modelling based on resting-state EEG data in the selection of patients for deep brain stimulation and potentially for other (non-invasive) interventions such as transcranial magnetic stimulation, transcranial direct-current stimulation, or vagal/median nerve stimulation Briand et al. 2020; Gosseries, Thibaut, et al. 2014. Particularly for MCS^*^ patients with higher recovery rates at the group level Thibaut, Panda, et al. 2021, the model would be valuable for predicting recovery. However, because the number of patients with MCS^*^ in this study was relatively small, they were not included in a separate group. We denoted them with different colors in the plots, which showed that the potential plasticity in the UWS group was mainly driven by MCS^*^ patients. Future studies using longitudinal data could assess whether this framework opens avenues for predicting therapeutic effects and spontaneous recovery in patients with DoC.

fMRI model-based approaches to better characterize DoC have shown that low-level states of consciousness are characterized by segregated and less connected network states, potentially caused by increased inhibition López-González et al. 2021; Luppi, Cabral, et al. 2023; Luppi, Mediano, et al. 2022. These approaches are increasingly being studied; however, their direct clinical relevance is limited because of the complexity of data acquisition and analysis. An important methodological advancement of the proposed approach is the induction of plasticity in the modelled resting-state EEG data. This could be relevant not only to DoC but also to other pathologies. EEG-based modelling studies on DoC are scarce Assadzadeh et al. 2023; N. D. Schiff, Nauvel, and Victor 2014, although we envision their clinical applicability to be more direct. For example, the analysis and (plasticity-induced) peak detection can be fully automated. Some reflections on the model-based approach are warranted. Our results may be biased by the choice of biophysical model, as this model for healthy conditions is tuned to generate a 1/f spectrum superimposed by alpha oscillations from corticothalamic resonance due to a delay in propagation between the thalamus and the cortex Peter A Robinson et al. 2001. However, previous work has demonstrated that there are three possible ways to generate alpha oscillations in this specific model, which include the corticothalamic circuitry, as well as an intracortical circuit Hindriks and Putten 2013. Finally, we focused on the synaptic plasticity of excitatory synapses. However, in light of the mesocircuit hypothesis, the role of inhibition should also be studied, suggesting that disinhibition of the thalamus may play an important role in inducing excitatory input from the thalamus. More detailed biophysical models that include the caudate and globus pallidus could be used to analyze the role of plasticity within the context of disinhibition in future studies. Finally, despite the existence of several other hypotheses on consciousness that could be applied in the context of DoC, such as integrated information theory Tononi 2008 and the global neuronal workspace Mashour et al. 2020, these cannot be used to make quantitative predictions for EEG spectra from patients with DoC based on underlying neuronal circuits but require extraction of other features from EEG data Melloni et al. 2023. Hence, we restricted our interpretation of our findings to the context of the mesocircuit hypothesis.

In conclusion, we demonstrated the advantage of biophysical modelling in individual subjects with DoC, showing that the recovery of the corticothalamic circuitry comes with the reappearance of physiological brain rhythms in some patients with DoC. Although these findings underscore the predictive ability of the mesocircuit hypothesis, they may also have potential clinical implications. Examples include the prediction of recovery in patients with DoC and aid in the selection of patients with sufficient residual brain capacity for (non-invasive) treatment. Future work is needed to verify whether the recovery of the excitatory corticothalamic circuitry results in good neurological recovery.

## 1 Methods and Materials

### Experimental design

We included 145 patients (58 females, mean age 40 years ± 17 years) and 30 healthy control subjects (controls; 14 females, mean age 43 years± 15). Patients were diagnosed with MCS (n=95), MCS^*^ (n=12), or UWS (n=38). The etiologies were traumatic brain injury in 76 patients, anoxia after cardiac arrest in 50 patients, mixed anoxia after cardiac arrest and traumatic brain injury in 7 patients, stroke/hemorrhage in 19 patients, and other etiologies such as metabolic encephalopathy or infection of the central nervous system in 3 patients. The average time between injury and hospitalization/scanning was on average 2.3 years, with a standard deviation of 3.5 years. This study was approved by the Ethics Committee of the University Hospital of Li`ege. All healthy participants and their legal surrogates provided written informed consent to participate in the study. The level of consciousness of the patients was assessed using the Coma Recovery Scale-Revised (CRS-R) J. T. Giacino, Kathleen Kalmar, and John Whyte 2004, which was repeated at least five times to minimize clinical misdiagnoses Wannez et al. 2017. The patients’ diagnoses were based on the best behavioral scores obtained over repeated CRS-R assessments during the week of hospitalization.

### EEG data acquisition and preprocessing

Part of the EEG data was used in previous studies Carri`ere et al. 2020; Mortaheb et al. 2019; Rizkallah et al. 2019; Thibaut, Panda, et al. 2021. Our preprocessing pipeline was in accordance with the recent guidelines and recommendations for EEG data analysis Pernet et al. 2020. Briefly, EEG recordings of 20–25 min were obtained from patients with DoC and healthy controls. The data were acquired using a high-density EEG system with a sampling frequency of 250 Hz. Data from some patients were acquired at a sampling frequency of 500 Hz. Data from these patients 418 were down-sampled at a sampling frequency of 250 Hz. An Electrical Geodesic Inc. EEG system with 256 channels was employed, and a saline solution was used. During data collection, patients were kept awake as much as possible. An examiner was present during the acquisition to ensure this. EEG data were imported into MATLAB version R2021a for further analysis, partly using FieldTrip Oostenveld et al. 2011. EEG data from the neck, forehead, and cheek channels were discarded because they can be easily contaminated by muscle artifacts, resulting in 150 channels that were used for further analysis. EEG data were segmented into epochs of two seconds, and epochs with signals that had an amplitude exceeding 100 *µ*V were rejected automatically. The remaining bad epochs were rejected via visual inspection (Jitka Annen). Data were referenced to a common average montage Thibaut, Panda, et al. 2021. The EEG data were further preprocessed using a zero-phase sixth-order Butterworth bandpass filter of 0.5–40 Hz. After preprocessing, we computed the power spectrum for each channel and averaged it across the channels. Only the scalp channels covering the parietal, temporal, and occipital areas were included to compute the grand average across the channels. Other channels, such as those covering the frontal areas and jaws, were excluded. Power spectra were computed using Welch’s method with windows of 5 s with 2.5 s overlap.

### FDG-PET data

Patients fasted for at least six hours prior to the FDG-PET procedure. FDG-PET data were acquired after intravenous injection of 150–300 MBq of FDG on a Philips Gemini TF PET-CT scanner. The PET data were spatially normalized and smoothed using a Gaussian filter with a full-width at half-maximum value of 14 mm. The data were recorded in a single 12-min emission frame after a 30-min uptake phase, and the images were corrected for attenuation. Two FDG-PET analyses were performed. First, a statistical evaluation of relatively preserved and hypometabolic regions compared to healthy volunteers was performed as described previously Panda, López-González, et al. 2023. The brain FDG-PET SUV of each patient with an unequivocal and reliable bedside diagnosis of UWS or MCS was visually inspected by three experts in the analyses of FDG-PET of DoC patients. The patients were blinded to their clinical diagnoses. Patients with clinical UWS were subsequently qualitatively classified as MCS^*^ Gosseries, Zasler, and Laureys 2014 based on the relative preservation of metabolism in the frontoparietal network Stender, Gosseries, et al. 2014. Second, for quantitative assessment of glucose metabolism, the FDG-PET metabolic index of the best-preserved hemisphere (MIBH) was calculated to approximate the cerebral metabolic rate of glucose at the single-subject level Stender, Mortensen, et al. 2016. As previously described Stender, Gosseries, et al. 2014, individual images were registered on a population-specific FDG-PET template. Images were segmented into the left and right cortices and extracerebral tissue. Cortical uptake was normalized based on the metabolism of the extracerebral tissue in healthy volunteers and scaled between 0 and 1, based on the mean activity of the extracerebral tissue. Finally, the MIBH was calculated as having the highest mean metabolism in both hemispheres.

### Corticothalamic mean-field model

We employed a corticothalamic mean-field model Peter A Robinson et al. 2001, which describes the aggregate activity of a neuronal population in terms of their firing activity *ϕ*_*a*_ and mean membrane potential *V*_*a*_ with a being either *e, i, r*, or *s*. The corticothalamic mean-field model encompasses two cortical populations (excitatory (*e*) and inhibitory (*i*)) and two thalamic populations (relay (*s*) and reticular (*r*)). The membrane potential of a population fluctuates *V*_*a*_(*t*) as a result of the incoming firing rate *ϕ*_*a*_(*t*) from other populations and/or itself according to R. Abeysuriya and P. Robinson 2016; P. Robinson et al. 2001; Peter A Robinson et al. 2001

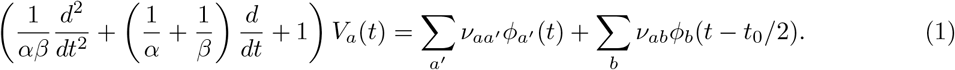

The effect of the presynaptic input on the postsynaptic membrane potential depends on the mean number of synapses between the presynaptic *b* and postsynaptic population *a* (modelled as synaptic strength *ν*_*ab*_) and on the closing and opening rate of synaptic channels characterized by the synaptic decay and rise constants (*α* and *β*). The average postsynaptic membrane potential is transformed at the cell bodies in a population, giving rise to the firing activity *Q*_*a*_(*t*)

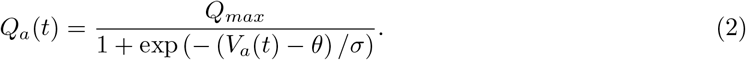

Here, *Q*_*max*_ refers to the maximum firing rate in *Hz, θ* is the mean firing threshold in *mV*, and *σ* is the standard deviation of this threshold. This process results in activity propagation in a closed loop between the thalamic and cortical populations, where the firing rate propagated between the thalamus and cortex is delayed by *t*_0_. The firing activity *Q*_*a*_(*t*) was temporally damped using the following expression

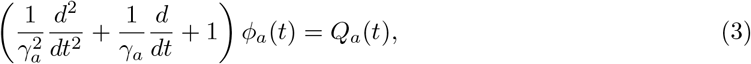

with *γ*_*a*_ being the temporal damping rate, based on *γ* = *v*_*a*_*/r*_*a*_, where *v*_*a*_ is the propagation velocity and *r*_*a*_ is the mean range of axons. For inhibitory, relay and reticular populations, *γ*_*a*_ *≈ ∞*, hence *ϕ*_*a*_(*t*) = *Q*_*a*_(*t*).

### Estimation of model parameters

#### Model power spectrum

Parameter estimation for nonlinear models remains challenging. Therefore, we transformed the nonlinear model into a linear model using linearization around a stable fixed point. Linearization is achieved by expressing the sigmoid function (Equation 2) that transforms *V*_*a*_(*t*) into *Q*_*a*_(*t*) as Taylor expansion and retaining only the term containing the first derivative (*ρ*_*a*_) evaluated at the fixed point. The corresponding details have been discussed previously R. Abeysuriya and P. Robinson 2016. Using the derivative (*ρ*_*a*_), we can express the synaptic strengths as gain parameters in the linear regime *G*_*ab*_ = *ρ*_*a*_*ν*_*ab*_, from which we can define loop parameters such as *G*_*aba*_ = *G*_*ab*_*G*_*ba*_. Accordingly, we transformed the linear system in the time domain to the frequency domain (with***k*** and *ω* being the wave vector and angular frequency) and derived the following transfer function with the firing rate of the excitatory population as output *ϕ*_*e*_(***k***, *ω*) and external signal*ϕ*_*n*_(***k***, *ω*) as input

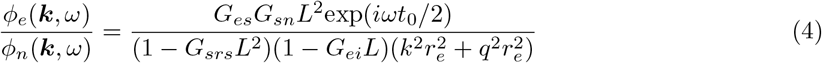

Here, *L* follows from the transformation of the second-order differential operator describing the synaptic response (Equation 1) in the Fourier domain, which can be interpreted as a low-pass filter depending on the synaptic parameters *α* and *β*

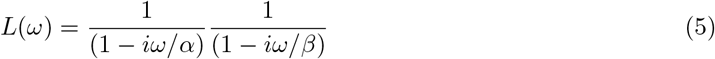

The EEG power spectrum can then be obtained by integrating over ***k***, where the cortex is approximated as a rectangular sheet due to its finite size, with a size of 0.5 m Timofeev et al. 2000. Using periodic boundary conditions, we derived the following power spectrum P. Robinson et al. 2001

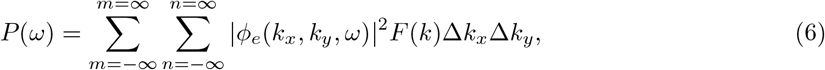

with 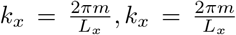,and 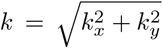. The filter function *F* (*k*) accounts for the low-pass filtering resulting from volume conduction by the skull, cerebrospinal fluid, and scalp Srinivasan, Nunez, and Silberstein 1998

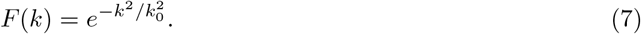

Here, *k*_0_ corresponds to the low-pass cutoff at *k*_0_ = 10*m*^*−*1^. This value was obtained in a previous study by using a spherical harmonic head transfer function Srinivasan, Nunez, and Silberstein 1998 Lastly, pericranial muscles activity resulted in electromyogram (EMG) artifacts in the EEG, necessitating our attention and correction; hence, *P*_*total*_(*ω*) = *P* (*ω*) + *P*_*EMG*_(*ω*).

#### Fitting model parameters

Similarly as in R. Abeysuriya and P. Robinson 2016 a selection of parameters was fixed to reduce parameter space, *Q*_*max*_ = 340*s*^*−*1^, *γ*_*e*_ = 116*s*^*−*1^, *θ* = 12.9*mV, σ* = 3.8*mV*. The parameter set *x* = [*G*_*ei*_, *G*_*ee*_, *G*_*ese*_, *G*_*esre*_, *G*_*srs*_, *α, β, t*_0_, *P*_*EMG*_] is estimated from EEG data by minimizing the error between the experimentally obtained power spectrum *P*_*exp*_ and model power spectrum *P*_*total*_(*x*) expressed as

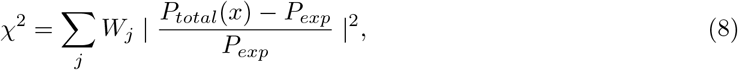

where *j* denotes the frequency bins. The weights *W*_*j*_ ensure equal weighting for every frequency decade and are proportional to 1*/f*. Because the parameter space was very large, we restricted the parameter values to neurophysiologically plausible values (see reference R. Abeysuriya and P. Robinson 2016 for values). The *χ*^2^ statistic is further transformed into a likelihood function as follows

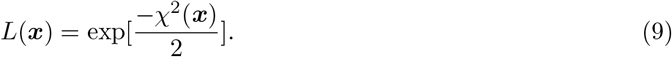

Hence, minimizing the error translates into maximizing the likelihood function. The Metropolis-536 Hastings algorithm was used to generate a probability distribution for each parameter using Markov Chain random walk Rosenthal et al. 2011. The details of the algorithm have been reported previously R. Abeysuriya and P. Robinson 2016. For every subject, we ran the Metropolis-Hastings algorithm to obtain model parameters for individual power spectra. The random walk was initialized using parameters obtained from a large database of healthy controls R. Abeysuriya, Rennie, and P. Robinson 2015. This initialization generally does not affect the final output but affects the convergence time. For every subsequent step in the random walk, the likelihood of this step was computed using Equation 13.

A new randomly proposed set is generated. The likelihood of this new set of parameters was computed using Equation 13. If these new parameters have a higher probability, this step is used to sample the probability distribution. Otherwise, a random number is drawn from a uniform distribution. If this random number is smaller than the ratio of the probability of the new parameters to that of the old parameters, the probability distribution sampling step is accepted. If this random number is larger than the ratio of the probability of the new parameters to that of the old parameters, the step is not accepted. This procedure was repeated several times until there were no iterative changes in the sampled probability distribution.

### Modelling synaptic plasticity

After estimating the subject-specific mean-field parameters, we performed simulations using a fully nonlinear model (Equations 1–3) to analyze potential routes for neurophysiological recovery in individual patients. In order to do this, we first transformed the subject-specific and estimated gain parameters *G*_*ab*_ to synaptic strength parameters *ν*_*ab*_ using the derivative of the sigmoid function *ρ*_*a*_ evaluated at the steady state *G*_*ab*_ = *ρ*_*a*_*ν*_*ab*_. The steady-state firing rate for each subject was obtained during the parameter estimation.

We initially explored two possible synaptic plasticity rules based on Hebbian and homeostatic plasticities Magee and Grienberger 2020. Hebbian plasticity refers to a positive-feedback-mediated form of plasticity in which synapses between presynaptic and postsynaptic neurons that are coincidently active are strengthened. Homeostatic plasticity refers to a negative-feedback-mediated form of plasticity, also known as synaptic scaling, which maintains the network activity at a desired set point. Both types of plasticity are thought to occur in neural systems Ho, Lee, and Martin 2011. We first evaluated the Hebbian plasticity using an existing implementation of our current mean-field model, also known as spike-timing-dependent plasticity Fung, Haber, and P. Robinson 2013 In addition, we implemented a general homeostatic plasticity rule, as used in previous studies Romesh G Abeysuriya et al. 2018; Hellyer et al. 2016

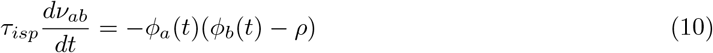

This Equation models the change in synaptic strength *ν*_*ab*_ between populations *a* and *b* as a function of their firing rate and a desired set point *ρ*, which will be set to the average firing rate from healthy control subjects that were included for this study. The synaptic time-scale parameter *τ*_*isp*_ is chosen exactly to be similar to that reported in previous literature and is set to *τ*_*isp*_ = 20 Romesh G Abeysuriya et al. 2018. Simulations of Equations 1–3 are executed, first, for the first 30 s without plasticity, succeeded by switching on plasticity for the remaining part of the simulations (10 min). First, we examined spike-timing-dependent plasticity. However, spike-timing-dependent plasticity in isolation leads to an unopposed strengthening of synapses, resulting in an unstable system. Next, we executed both spiketiming-dependent plasticity and homeostatic plasticity but noticed that stability only occurred if the contribution of homeostatic plasticity was very large and that of spike-timing-dependent plasticity was negligible. Hence, in the results presented in our manuscript, we reported only homeostatic plasticity. Simulations of the model are performed by solving Equations 1–3 and 14 using an Euler-Maruyama solver with a sufficiently small time step (1×10-4) using in-house Matlab scripts (R2021a). The Laplace operator in Equation 3 is set to zero to run the model as a neural mass model instead of a neural field model.

### Statistics

We test for significant differences for parameters (*G*_*ei*_, *G*_*ee*_, *G*_*ese*_, *G*_*esre*_, *G*_*srs*_, *α, β*, and *t*_0_) between groups using the Wilcoxon rank sum test. The false discovery rate was subsequently used to correct for multiple tests Benjamini and Hochberg 1995. We quantified the similarity between the power spectra for different groups using the Kullback-Leibler divergence Van Erven and Harremos 2014. Summary statistics from the power spectra, such as the slope, peak frequency, and peak power, were extracted using the FOOOF algorithm Donoghue et al. 2020. We used the following settings for the FOOOF algorithm (maximum number of peaks = 2, minimum peak height 0.3 (units of power), peak width limits 1-14 (Hz), aperiodic mode = “knee”). We tested the null hypothesis that the presence of a spectral peak was equal for MCS and UWS using the chi-square test. Finally, we tested the relationships between these summary statistics and the metabolic indices using Pearson’s correlation coefficients.

## Data and materials availability

Data and materials availability: The code for the parameter estimation method can be found at https://github.com/BrainDynamicsUSYD/braintrak, and includes an explanation of the method on the wiki page https://github.com/BrainDynamicsUSYD/braintrak/wiki. Data cannot be shared on open-sharing platforms due to the lack of informed consent from patient representatives. Further ethical approval is required for data sharing.

## Author Contributions

Conceptualization PKBT, JA, GD, SL, AT, OG; Methodology PKBT, RA, AT, JA, OG Investigation: PKBT, RA, JA, PN, MMV, GL, RP; Visualization: JA, PKBT, RA, RP; Supervision JA, GD, SL Writing—original draft PKBT, JA; Writing—review and editing: PKBT, RA, RP, PN, MMV, GL. OG, AT, SL, GD, JA

## 2 Acknowledgments

We thank the patients and their families who agreed to participate in this study. The authors also thank the staff of the Nuclear Medicine Department at the University Hospital of Liege, especially Roland Hustinx and Claire Bernard. We are highly grateful to the members of the Li`ege Coma Science Group/Centre du Cerveau for their assistance in the clinical evaluations, especially Cecile Carette, and the clinicians from the Intensive Care Unit of the University Hospital of Liege, especially Didier Ledoux, Paul Massion, and Gaelle Tronconi. PT was supported by the EMBO New Venture Fellowship 9139, EAN Research Experience Fellowship, and Jan-Meerwaldt Fellowship awarded to PKBT. This study was funded by the University and University Hospital of Liege, the Belgian National Funds for Scientific Research (FRS-FNRS), the European Union’s Horizon 2020 Framework Programme for Research and Innovation under Specific Grant Agreement No. 945539 (Human Brain Project SGA3), the FNRS MIS project (F.4521.23), and the FNRS PDR project (T.0134.21). AS is a postdoctoral fellow, OG and AT are research associates, and SL is the research director at the F.R.S.-FNRS. S.L. received funding from the European Union’s Horizon 2020 Framework Programme for Research and Innovation under Specific Grant Agreement 768 Nos. 785907 (Human Brain Project SGA2) and 945539 (Human Brain Project SGA3). The study was further supported by the University and University Hospital of Liege, the Belgian National Funds for Scientific Research (FRS-FNRS), the European Space Agency (ESA), and the Belgian Federal Science Policy Office (BELSPO) in the framework of the PRODEX Programme, FNRS PDR project (T.0134.21), the FNRS MIS project (F.4521.23) the University of Li`ege Conseil Sectoriel de la Recherche, “Fondazione Europea di Ricerca Biomedica”, the BIAL Foundation, the Mind Science Foundation and the European Commission, the fund Generet, the King Baudouin Foundation, AstraZeneca foundation, Leon Fredericq foundation, the National Natural Science Foundation of China (Joint Research Project 81471100), and the European Foundation of Biomedical Research FERB Onlus.

## Notes

### Competing Interest Statement

The authors have declared no competing interest.

